# Beaver population decline on Michipicoten Island, Ontario leads to satellite-measured surface water area reductions

**DOI:** 10.64898/2026.02.20.707078

**Authors:** Robert H. Fraser, Ian Olthof, Ashley McLaren, Brent Patterson

## Abstract

The North American beaver (*Castor canadensis*) is an ecosystem engineer that strongly influences stream hydrology and ecosystems by constructing dams and canals. Previous research has shown that changes in the extent of beaver ponds and wetlands mapped using aerial photographs can serve as a proxy indicator of shifting regional abundance of beavers. In this study we investigated the use of freely available optical satellite data to measure changes in beaver pond surface water area on the 184 km^2^ Michipicoten Island in Lake Superior (Ontario, Canada) after a large decline in the beaver population that followed the arrival of grey wolves (*Canis lupus*). Inter-annual variability in pond extents was measured using sub-pixel mapping methods applied to the 30 m resolution Landsat (1985-2023) and 10 m Sentinel-2 (2016-2023) satellite records. After a > 90% decline in the number of surveyed beaver colonies between 2015-2018, beaver pond surface water area was reduced by 38-42% for ponds < 0.5 ha and by 48% for ponds < 0.1 ha by 2023. While these recent ponding reductions occurred during a period of above average precipitation, two previous smaller reductions were associated with low precipitation, water balance index, and Lake Superior water levels, suggesting that they were caused by drought and not beaver colony declines. While further testing is warranted, our results show that satellite-mapped changes in beaver ponds can provide a cost-effective metric for assessing large-scale population trends in the boreal zone.

## INTRODUCTION

The North American beaver *(Castor canadensis)* is broadly distributed in aquatic habitats across North America, where populations have been recovering since the last century after being decimated by overexploitation (Rosell and Campbell-Palmer 2022). Where beavers build dams across stream channels and floodplain valleys, they have a major impact on ecosystems by retaining water and sediment, removing vegetation for food and building material, and creating unique, biodiverse habitats (Larsen et al. 2021, Grudzinski et al. 2022, Fairfax and Westbrook 2024). Beavers also buffer against drought, attenuate flood flows, and provide fire refugia, thus representing a promising nature-based method of reducing the effects of climate change (Fairfax and Westbrook 2024). To better assess changing beaver distributions and densities, and these wide range of impacts, efficient methods are needed for monitoring the spatial and temporal distribution of beavers.

Beaver populations have traditionally been surveyed by detecting recent signs of beaver colony activity, such as fresh mud and peeled sticks placed on lodges or dams (Rosell and Campbell-Palmer 2022,). In colder climates where beaver ponds freeze during winter, each colony normally constructs a food cache in autumn near the lodge by submerging forage under a raft of less desirable tree species (Slough 1978). Surveying these autumn food caches using airborne surveys (Bergerud and Miller 1977, Ribic et al. 2017, Johnson-Bice et al. 2021, Thompson et al. 2022) or ground observations (Fryxell 2001, Hood and Bayley 2008) is therefore a common approach to census beaver colonies.

Another, indirect method for measuring beaver population fluctuations is to track the presence of ponds and wetlands that are created by their maintained dams. This usually involves interpreting beaver ponds in an irregular time series of aerial photographs taken over multiple decades (Broschart et al. 1989, Johnston and Naiman 1990, Meentemeyer and Butler 1995, Snodgrass 1997, Cunningham et al. 2006, Martell et al. 2006, Hood and Bayley 2008, Johnston and Windels 2015, Martin et al. 2015). More recently, high-resolution (∼ 1 m) satellite images have been used to map beaver ponds (Malison et al. 2014), including their expansion into the Alaskan tundra (Tape et al. 2022). While long-term variation in beaver ponds was associated with changing regional abundance of beaver colonies in these studies, this approach can be confounded by beavers damming the most favourable sites first that create larger ponds (Johnston and Naiman 1990, Cunningham et al. 2006), and lags in pond drainage after they are abandoned by beavers (Woo and Waddington 1990, Johnston and Windels 2015, Johnson-Bice et al. 2022).

Coarser resolution satellite mapping using temporally consistent and freely available 30 m Landsat images from 1984 to present has also been used to track beaver ponds (Townsend and Butler 1996, Tape et al. 2018, Jones et al. 2021, Fraser et al. 2024, Fraser et al. 2025). After observing that existing binary Landsat surface water products are insufficient for this application (Fig. S1), Fraser et al. (2024) demonstrated the use of sub-pixel surface water mapping (Olthof and Fraser 2024) to track inter-annual variation in beaver ponds in Manitoba’s sub-arctic coastal zone. A large (> 80%) multi-year reduction measured in the area and number of beaver ponds was inferred to be associated with a beaver population decline. However, aside from observing new dams in high-resolution imagery for a selection of recovering ponds, this linkage could not be directly demonstrated without historical beaver censuses.

In this study, we applied this Landsat sub-pixel mapping approach to investigate changes in surface water area resulting from a recent documented beaver colony decline on Michipicoten Island in the Boreal Shield zone of Ontario, Canada (McLaren et al. 2022). We hypothesized that, after a time lag, the total surface water area of beaver-created ponds would decrease after ponds were abandoned by beavers and their dams became unmaintained. We also examined longer-term, 38-year changes in Landsat-derived beaver pond area to assess their relationships with climate data and compared Landsat estimates to those from higher resolution Sentinel-2 (10 m) and WorldView/GeoEye (∼1 m) satellite data. The broader goal of this research was to develop and test satellite remote sensing methods for tracking indicators of beaver population changes in remote landscapes where conventional monitoring is challenging or prohibitively expensive.

## MATERIALS AND METHODS

### Study Area

Michipicoten Island covers 184 km^2^ and lies within the Ontario, Canada portion of Lake Superior (Fig. 1). The island is the third largest in Lake Superior and was designated a natural environment provincial park in 1985. Its rugged uplands where most of the abundant beaver colonies reside contain east to west ridges with gentle southern slopes and sleeper northern slopes. The main streams follow the east-west valleys, while smaller tributary streams and creeks drain north-south along steeper slopes. Michipicoten Island is covered mainly by a mixture of southern boreal needleleaf forest (balsam fir and spruce) and northern Great Lakes-St. Lawrence broadleaf forest (white birch and sugar maple). Larger mammals on the island have included woodland caribou (*Rangifer tarandus caribou*), North American beaver, and red fox (*Vulpes vulpes*) (Ontario Parks 2004). The beaver population thrived due to a lack of natural predators of adult beavers until 3-4 grey wolves (*Canis lupus*) came to the island on an ice bridge in the winter of 2013-2014, growing to a population of 20 wolves by winter 2017-2018 (Patterson et al. unpublished data). A diminished and heavily preyed-upon caribou population was moved from the island in 2018 and eight of the remaining wolves were translocated to Isle Royale on Lake Superior in 2019. Following this translocation at least two wolves remained on the island through March 2022 (Patterson et al. unpublished data). A more detailed description of Michipicoten Island and the history of its beaver population that has been unharvested since the mid-1980s is provided in McLaren et al. (2022).

**Figure 1.**
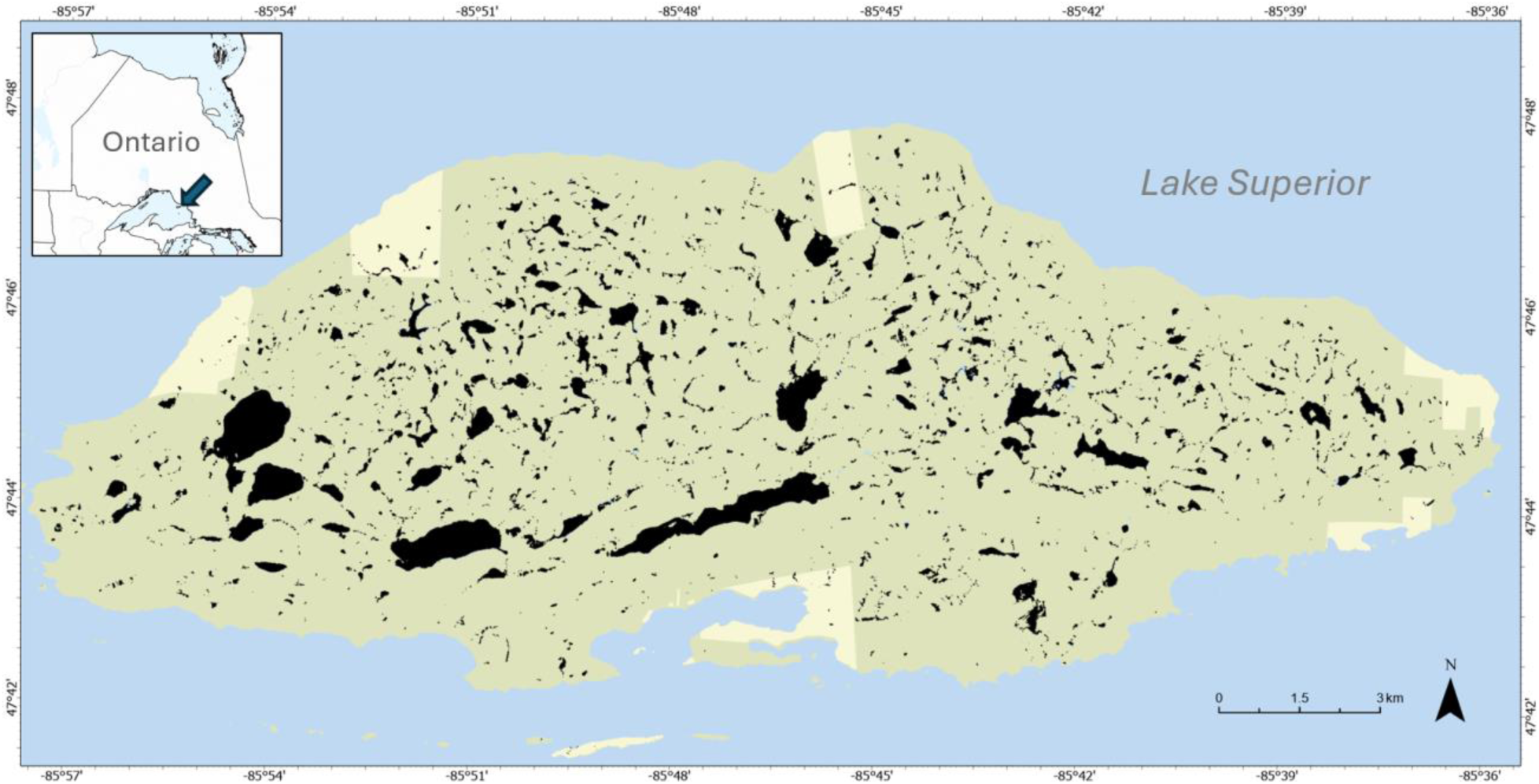
Michipicoten Island lying within the Ontario, Canada portion of Lake Superior. The maximum 2016-2023 surface water extent of 2325 unique water bodies mapped using Sentinel-2 satellite data is shown in black (Background: Esri World Street Map, Sources: Province of Ontario, Esri Canada, Esri, TomTom, Garmin, SafeGraph, FAO, METI/NASA, USGS, EPA, NRCan, Parks Canada).

### Landsat satellite mapping of surface water area changes

We mapped the surface water area on Michipicoten Island annually for 1985-2023 using Landsat data and a sub-pixel mapping method (Olthof and Fraser 2024) that was previously applied to map long-term changes in beaver ponds within > 4500 km^2^ study regions in northern Manitoba (Fraser et al. 2024) and northwestern Ontario (Fraser et al. 2025). The method estimates the percentage surface water cover (*f_water_*) within each 30 m pixel using linear unmixing of measured near-infrared (NIR) pixel reflectance (*p_NIR_*) that is a linear combination of the reflectance of water and land endmembers (Eq. 1) (Settle and Drake 1993). In this implementation, water endmember reflectance (*E_waterλ_*) was derived from the median reflectance of all permanent surface water bodies and a spatially variable land endmember reflectance (*E_landλ_*) was adaptively sampled from background land pixels (Olthof and Fraser 2024).

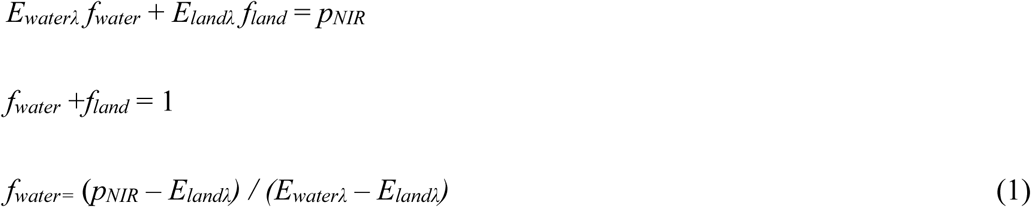

Annual fractional surface water area maps were created for Michipicoten Island by applying this technique to Landsat NIR reflectance composites from 1985-2023. The composites represented rolling, three-year median surface reflectance values from June 1 – September 30 cloud-free observations. They were generated in Google Earth Engine using all Collection 2 Tier 1 Level 2 Landsat images that had < 50% cloud cover (Olthof and Fraser 2024).

Needleleaf tree canopies and their cast shadows can produce erroneous small water fraction predictions because of their low NIR reflectance (Olthof and Fraser 2024). To address this issue, we removed water pixels containing less than 15% maximum surface water and derived a mask representing needleleaf forest on Michipicoten Island. This mask was created by manually thresholding the red reflectance channel from a clear-sky, late-winter (20230410) Sentinel-2 image. In this image, the red channel presents a bright signal over snow-covered water bodies and open areas (e.g. previously flooded beaver meadows) and a contrasting dark signal over needle-leaf forest (Fig. S2). The maximum 1985-2023 surface water percentage was calculated for the resulting unmasked water pixels, which were then spatially clustered to form unique multitemporal water objects following Fraser et al. (2024). These objects, the vast majority of which are beaver-created disturbance patches (Johnston and Naiman 1990), were then visually confirmed as being either current water bodies or drained ponds using 0.5 m resolution pan-sharpened WorldView imagery. This resulted in the removal of 4% of falsely mapped water objects that usually corresponded to topographic shadows.

### Sentinel-2 satellite mapping of surface water area changes

Surface water was also mapped annually for 2016-2023 at a higher 10 m spatial resolution using the Sentinel-2 satellite record. The spectral unmixing method we used to map Landsat fractional water requires a long-term surface water product to identify stable endmember samples (Olthof and Fraser 2024). Since such a product is not available for Sentinel-2, we applied a different linear unmixing approach. This method uses a reflectance frequency histogram to define reflectance breakpoint values representing the boundaries of pure water and pure land pixels (Olthof et al. 2015). The water fraction of mixed pixels (*f_mwater_*) lying within these pure water (WL) and land (LL) reflectance limits is then determined by linear interpolation (Eq. 2). For Michipicoten, manual inspection of the NIR reflectance histogram from a July 25, 2018 Sentinel-2 Level 1C image (Table S1) was used to derive breakpoint reflectance values of 0.068 for water (WL) and 0.183 for land (LL). A Level 1C image representing top-of-atmosphere reflectance was used instead of Level 2A surface reflectance to be compatible with the Level 1C harmonized Sentinel-2 collection available in Google Earth Engine that was used to derive 2016-2024 surface water fractions.

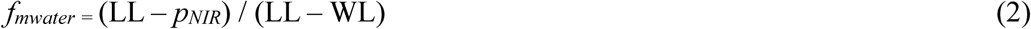

Annual Sentinel-2 NIR reflectance values for 2016-2023 were also derived in Google Earth Engine based on rolling three-year median compositing. All images from June 10 – September 20 were used and a Cloud Score+ (Pasquarella et al., 2023) threshold of 0.80 was applied to remove pixels containing cloud and shadow. Percent water fraction for each year was then derived using these median NIR values and the histogram breakpoints. As in the Landsat processing, the maximum multiyear surface water was calculated, masked using the needle-leaf forest classification, and then grouped into unique, spatially connected water objects for computing annual surface water areas. Three percent of the Sentinel-2 water objects identified as falsely mapped water were removed. We also produced surface water maps using four cloud-free Level 1C Sentinel-2 images from 20160705, 20180725, 20230818, and 20240718 (Table S1) covering Michipicoten Island to compare single-date predictions to those derived from the three-year composites.

### Validation of Landsat and Sentinel-2 surface water estimates

The Landsat and Sentinel-2 surface water mapping methods were assessed over Michipicoten Island using a higher resolution (1.3 m) surface water map derived from a commercial WorldView-3 satellite image (July 18, 2018, catalogue ID: 104001003F552400) that covered the central half of the island. To create this map, the image was orthorectified using PCI Geomatics Catalyst software and the NIR channel manually thresholded to produce a surface water extent consistent with one interpreted from a pan-sharpened (0.3 m) RGB version of the image (Fraser et al. 2024). This yielded 5,331 high-resolution water bodies after removing noise due to topographic shadows. The WorldView-3 map was used to assess Landsat and Sentinel-2 water bodies that were delineated over the same coverage area based on the methods described above. For this, single-date Landsat and Sentinel-2 images from July 25, 2018 were used rather than the median reflectance composites to most closely match hydrologic conditions corresponding to the high-resolution image date. These 10 m and 30 m resolution water bodies were spatially intersected with the WorldView-3 water bodies to facilitate a comparison of their surface water areas using linear regression.

A validation of two-date Sentinel-2 surface water changes was also conducted by deriving a more recent high-resolution surface water map from a July 4, 2024 orthorectified commercial GeoEye-1 image (catalogue ID: 105001003C20A30). For this validation, high-resolution surface water changes from July 18, 2018 to July 4, 2024 were compared to Sentinel-2 surface water changes derived from clear-sky images available for July 25, 2018 and July 18, 2024. Change estimates for 933 water bodies were compared using regression analysis from areas where there was common cloud-free coverage in the high-resolution images.

### Comparing changes in beaver colony presence and surface water area

The number of active beaver colonies on Michipicoten Island was determined for 2015 and 2017-2019 by recording the annual presence of autumn food caches. Instead of using the conventional approach of visually surveying caches from aircraft, a new image-interpretation method was developed to account for the island’s exceptionally high beaver density (McLaren et al. 2022). This involved capturing 7-8.5 cm resolution overlapping photographs from systematic airborne flight lines and creating image orthomosaics covering the island using structure-from-motion photogrammetry methods. Comprehensive, manual interpretation and digitizing of the annual orthomosaics revealed that the number of beaver food caches on Michipicoten Island declined by > 90%, from 1,119 (6.1 / km^2^) in 2015 to 67 (0.4 / km^2^) in 2018 (McLaren et al. 2022).

We did not attempt to match each beaver cache location with the corresponding water bodies maintained by a colony to investigate how these beaver colony reductions impacted surface water area for several reasons. First, colony territories frequently comprise several adjacent dams and ponds along a stream reach (Johnston and Naiman 1990), which would make it difficult to correctly attribute ponds to cache locations when colonies occurred at very high density as on Michipicoten Island. Second, artifacts and co-registration issues in portions of the annual orthomosaics prevented consistent tracking of individual cache status but permitted island-wide censusing of cache numbers (McLaren et al. 2022). Finally, we assumed that the surface water area changes caused by a > 90% reduction in colonies would dominate the change signal observed in beaver ponds when aggregated across Michipicoten Island.

Therefore, instead of attempting to attribute the changes in individual beaver ponds to the presence of a colony that created them, we summed the annual surface water area on the island according to different size classes: (a) the largest water bodies (> 30 ha) that were lakes containing no beaver dams adjacent to their outlets and were minimally impacted by beavers (n=7) (McLaren et al. 2022), (b) all other water bodies (< 30 ha), which were consistently beaver-influenced as evidenced primarily by dams but also lodges and canals visible in high-resolution (0.5 m) satellite imagery (Fig. S3), (c) the subset of these beaver ponds < 1 ha, and (d) the subset of beaver ponds < 0.5 ha. The finer resolution of Sentinel-2 permitted analyzing an additional beaver pond size class of < 0.1 ha.

### Investigating climate influences on long-term beaver ponded area

Although the major focus of this study was detecting surface water area changes resulting from a recently declining beaver population, we also investigated if there were long-term (1985-2023) relationships between annual Landsat beaver pond area and three water-related climate variables, which may have also influenced any detected trends. Because no long-term climate stations are located near the study area, annual precipitation for central Michipicoten Island was derived from the ERA5-Land reanalysis dataset that combines model data with observations into a globally consistent dataset from 1950 to present at a 9 km native resolution (Munoz Sabater 2019). We analyzed a running three-year average of annual precipitation to be compatible with the three-year median satellite reflectance data used to compute beaver pond area. As in Fairfax and Small (2018), we also examined a drought index called the Standardised Precipitation-Evapotranspiration Index (SPEI) that is based on rainfall and potential evapotranspiration (Vicente-Serrano et al. 2010). SPEI provides a standardized measure of water balance where positive values represent wetter conditions and negative values indicate drier conditions. Monthly SPEI estimates based on 0.5-degree gridded Climate Research Unit precipitation and temperature data were obtained over the study region from the Global SPEI database (Beguería et al. 2010). We used average annual summer (June-September) SPEI values calculated on a 36-month time scale to again match the temporal scale of the Landsat data. Finally, annual average June-September Lake Superior water levels (Great Lakes Coordinating Committee 2024) provided a long-term regional measure of water gains from lake precipitation and basin runoff, and losses from lake evaporation (Assel et al. 2004). Pearson correlations and inspection of time series plots were used to characterize the relationships between beaver pond area and these climate-related variables.

## RESULTS

### Annual satellite-based mapping of Michipicoten water bodies

A total of 840 long-term water bodies were delineated on Michipicoten Island using Landsat sub-pixel surface water mapping. The Landsat inter-annual variability in surface water area for the four size classes is shown in Fig. 2a. The largest seven water bodies, representing lakes having little or no impact from beavers, showed the smallest amount of variability and a 2.7% reduction in surface water area (i.e., ponding) during 2017-2023 after the rapid decline in active beaver colonies that began after 2015. Water bodies smaller than this that are beaver influenced (n = 833) demonstrated much larger inter-annual variability and a 27% reduction during 2017-2023, which is the largest multiyear decrease observed during the 1985-2023 analysis period. Progressively larger percentage reductions during 2017-2023 occurred in smaller pond classes of < 1 ha (−40%) and < 0.5 ha (−42%). Additional periods of ponding reductions were observed from 1986-1992 (14%) and 2001-2012 (15%).

**Figure 2.**
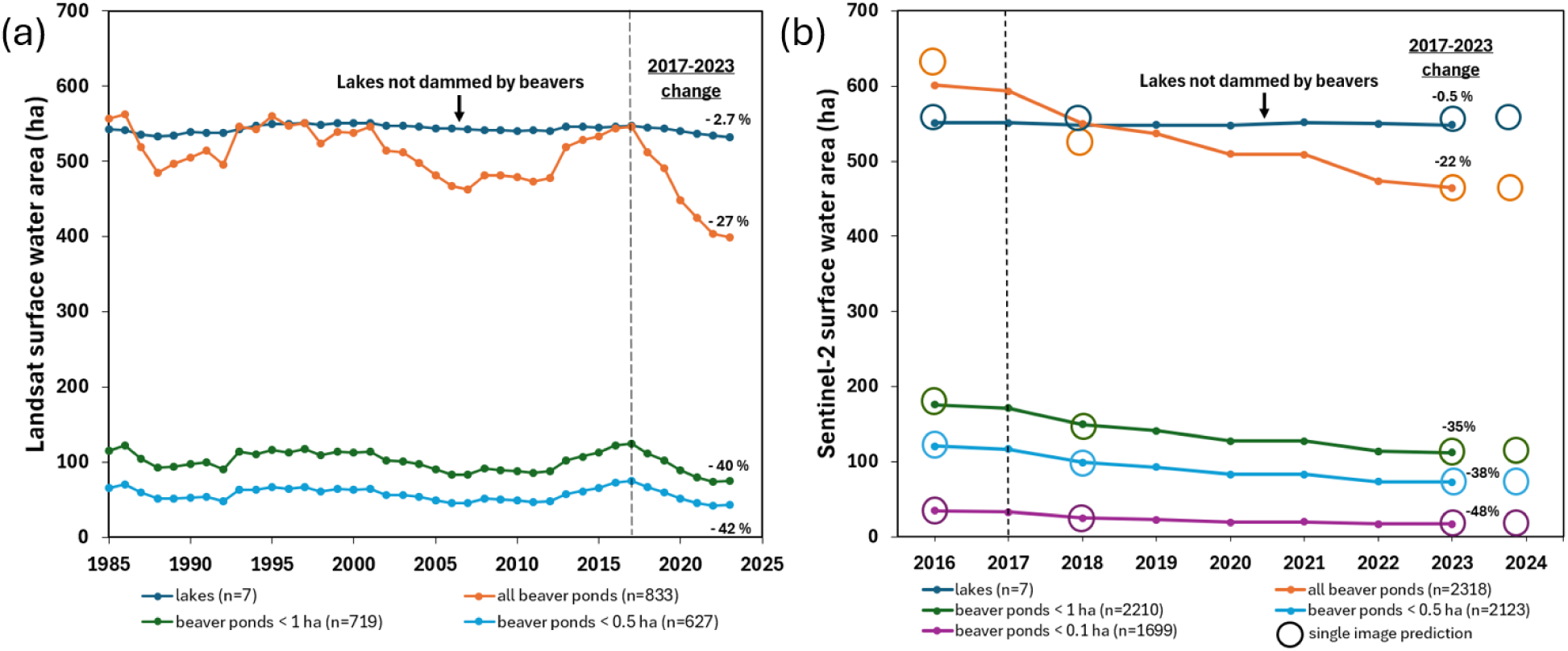
Change in Michipicoten surface water area by water object size class, mapped for 1985-2023 using Landsat data (a) and for 2016-2024 using Sentinel-2 data (b).

Sub-pixel water mapping using Sentinel-2 data delineated 2325 unique water bodies on Michipicoten Island (Fig. 1). While the higher resolution Sentinel-2 data captured a ∼ 10% larger area of beaver ponds, the 2017-2023 percentage declines (Fig. 2b) were similar to those measured using Landsat. In the smallest size class (< 0.1 ha or ∼ one Landsat pixel), Sentinel-2 showed a 48% surface water decline following the large drop in the number of active beaver colonies. The four, single-date Sentinel-2 surface water estimates (circles in Fig. 2b) closely corresponded to the median composite estimates, and the estimates for July 4, 2024 suggest that beaver pond area had stabilized at that time but not yet begun to recover to pre-2017 levels. Animations are provided that show changes in sub-pixel surface water mapped on the island between 2015-2017 and 2021-2023 using both Landsat (Supplementary Video 1) and Sentinel-2 data (Supplementary Video 2).

### Validation of Landsat and Sentinel-2 surface water

The Landsat sub-pixel water map for July 25, 2018 intersected with only 1475 / 5331 (28%) of the WorldView-3 water objects mapped for July 18, 2018. However, these objects accounted for 5,348,679 / 5,903,665 ha (91%) of the WorldView-3 surface water area. By contrast, the Sentinel-2 map for July 25, 2018 captured 2705 / 5331 (51%) of the WorldView-3 objects that represented 5,730,379 / 5,903,665 ha (97%) of the WorldView-3 water area.

Fig. 3 presents the relationships and linear regression coefficients between the areas of WorldView-3 water objects < 10,000 m^2^ (1 ha) and coarser resolution Landsat (Fig. 3a) and Sentinel-2 (Fig. 3b) objects that were captured using subpixel mapping. In these plots, WorldView-3 objects were merged when more than one intersected the same coarse-resolution water object, and vice-versa. Both Landsat and Sentinel-2 demonstrated a strong linear fit with WorldView-3 pond size, with overpredicted outliers in some cases being caused by the coarser resolution sensors mapping dark, wet sediment as water. For the subset of smaller WorldView-3 water objects < 2,500 m^2^ (0.25 ha), the regression fit using Landsat (Fig. 3c) degraded markedly compared to the Sentinel-2 fit (Fig. 3d). High-resolution surface water area changes from 2016-2024 are plotted in Fig. 4 against corresponding Sentinel-2 surface water changes for 933 water bodies. We found that the Sentinel-2 changes could explain 65% of the variation in water body changes mapped at higher resolution. When only more dynamic water bodies are considered that corresponded to Sentinel-2 ‘changes > 30% (n=673), the amount of variation explained increased to 78%.

**Figure 3.**
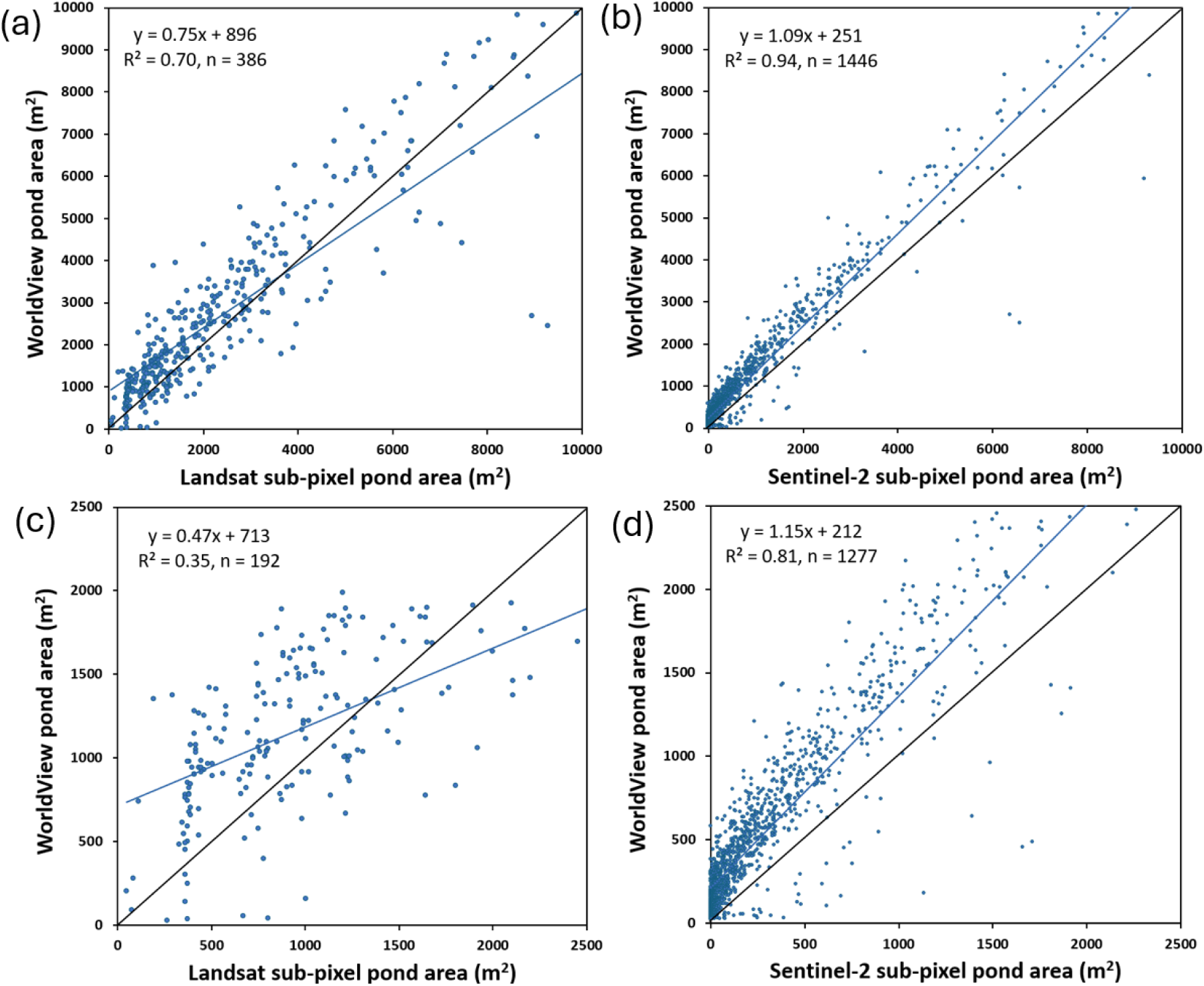
Comparison of intersecting water body areas mapped at high resolution using a WorldView-3 image with areas derived using medium resolution Landsat and Sentinel-2 data. WorldView ponds < 10000 m^2^ (1 ha) are shown in (a-b) and ponds < 2500 m^2^ are shown in (c-d). Linear least-squares regression equations, coefficients of determination (R), sample sizes (n), and 1:1 relationship lines (black) are included on the plots. All relationships are significant at p < 0.05.

**Figure 4.**
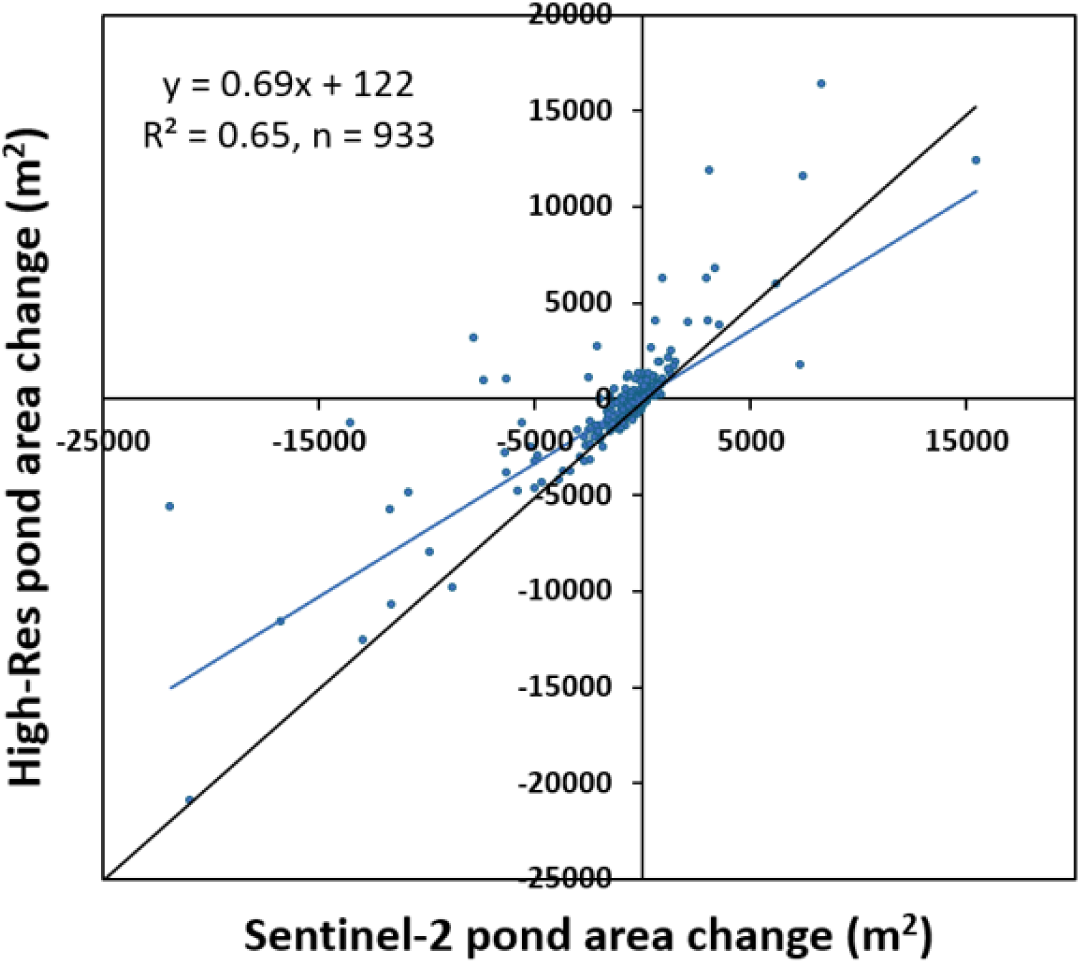
Comparison of water body changes mapped at high resolution (WorldView-3 and GeoEye-1) and medium resolution (Sentinel-2).

Fig. 5 and Fig. S4 present two-date comparisons between the high-resolution surface water mapping (a-d) and the coarser resolution Sentinel-2 (e-f) and Landsat (g-h) mapping. These highlight the ability for Sentinel-2 mapping to detect smaller ponds that are missed using Landsat and more precisely delineate the boundaries and areas of water bodies, as is indicated in Fig. 3. They also show that aquatic vegetation in beaver ponds results in surface water omission errors.

**Figure 5.**
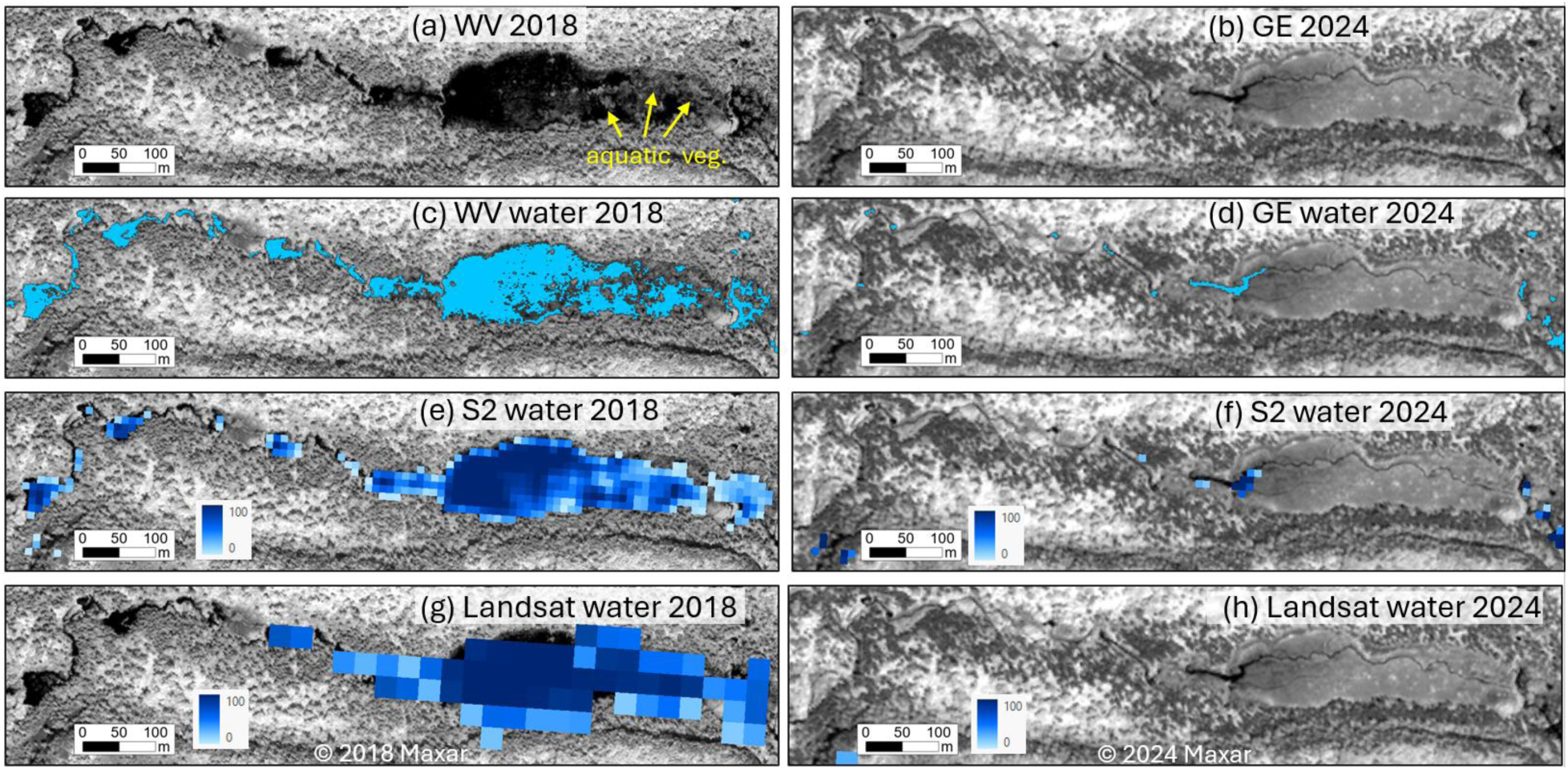
Example of 2018-2024 beaver pond changes mapped using high resolution WorldView-3 (a,c) and GeoEye-1 satellite images (b,d) and medium resolution Sentinel-2 (e-f) and Landsat (g-h) data. Surface water omission errors in 2018 can be observed at locations where there is a cover of floating aquatic vegetation.

### Relationship of long-term beaver pond surface water area to climate variables

The relationships between annual Landsat-derived beaver pond area and the climate variables representing water availability (Lake Superior water levels, annual precipitation, SPEI precipitation-evapotranspiration index) are shown in Fig. 6. All three variables demonstrated decreases during the first two ponding reductions in 1986-1992 and 2001-2012 and had moderate correlations (R^2^ = 0.25-0.55, p < 0.05) with ponded area from 1985-2017. During the larger 2001-2012 ponding decline, Lake Superior had its lowest annual water level since 1926 and ERA5 precipitation decreased to its smallest amount since 1951. After 2017, all three measures of wetness remained above their long-term averages and became disconnected from a steeply declining beaver pond area. When the full 1985-2023 time series was considered, the correlation between these variables and ponding became weak (R^2^ = 0.21, p < 0.05, Fig. 6b) or non-significant (R^2^ = 0.01-0.04, p > 0.05, Fig. 6a,c).

**Figure 6.**
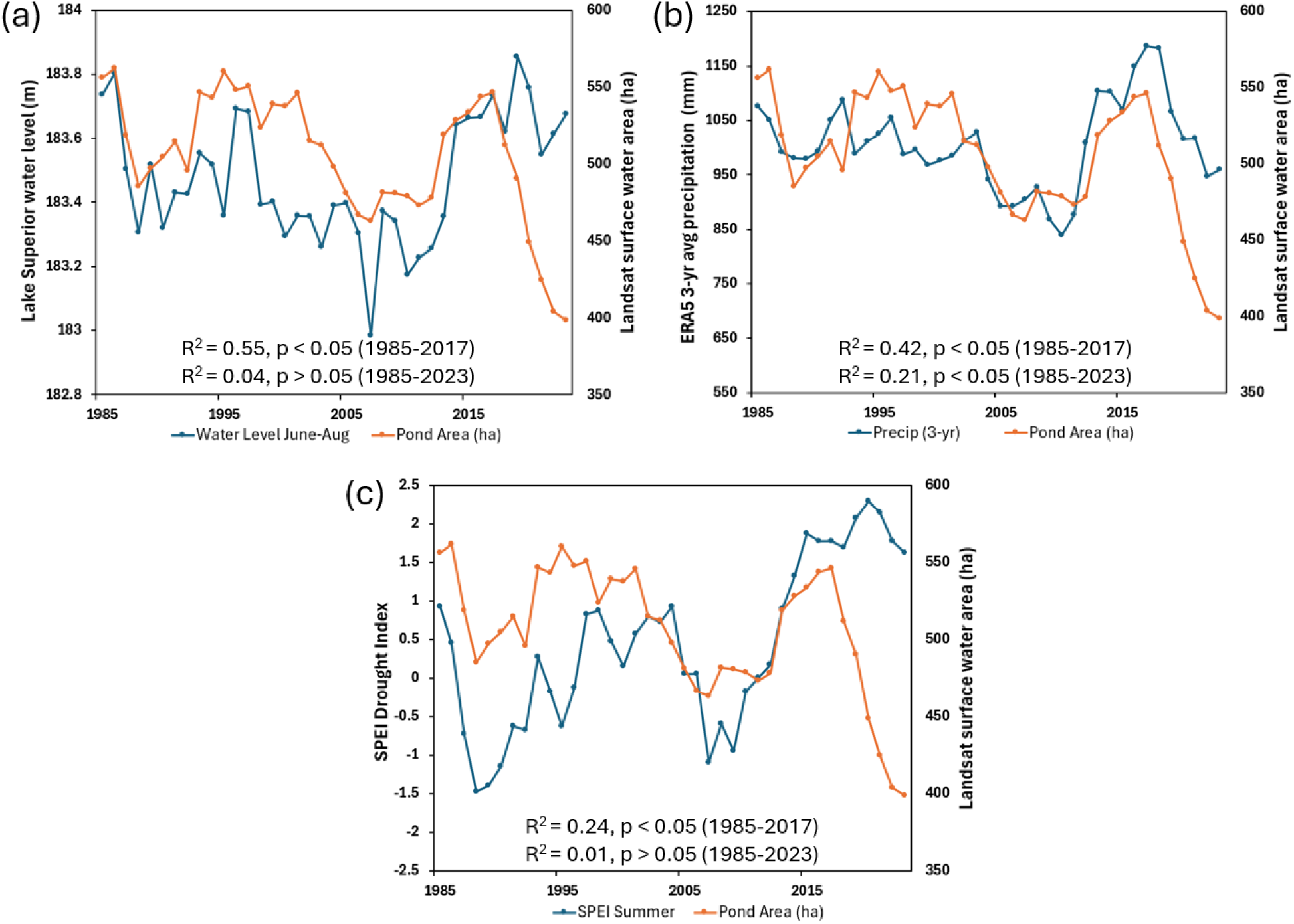
1985-2023 temporal associations between total Landsat-derived beaver pond area (833 beaver-engineered water bodies) on Michipicoten Island (orange) and (a) Lake Superior summer water levels, (b) ERA5-Land three-year average annual precipitation, and (c) summer Standardized Precipitation-Evapotranspiration Index (SPEI) where higher values represent relatively wetter conditions (blue).

## DISCUSSION

Our results demonstrate that freely available, medium resolution satellite data can be used to measure reductions in beaver ponded area associated with a regional decline in the number of beaver colonies. On Michipicoten Island, ponded area measured using both Landsat and Sentinel-2 data showed a large decrease from 2017-2023 that continued after 2018 when the smallest number of active colonies had been recorded. This lagged response indicates that beaver pond reductions were a gradual process on the island after colony abandonment, as unmaintained dams may slowly degrade and lose their ability to impound water (Woo and Waddington 1990). The fact that there was a > 90% decline in colonies but a < 50% reduction in beaver ponded area indicates that many ponds were resilient to becoming fully drained after their dams became unmaintained (Johnston and Windels 2015, Johnson-Bice et al. 2022, Fraser et al. 2024).

It is uncertain why smaller beaver pond size classes experienced progressively larger proportional decreases (Fig. 2). This could be related to surviving beaver colonies inhabiting larger ponds because of the increased protection they provide against wolf predation (Gable et al. 2018), or the greater availability of aquatic vegetation in larger, mature ponds (Ray et al. 2001) that would offer an alternative to broadleaf tree forage (Milligan and Humphries 2010) with a new predation risk over land. It is also likely that beavers had a smaller influence on the total surface water area of many larger ponds on Michipicoten Island that were enlarged but not originally created by beaver engineering.

The 10 m resolution Sentinel-2 data demonstrated a greater ability to detect smaller beaver ponds and more precisely measure their areas compared to Landsat. However, since the majority of beaver pond area loss was attributable to ponds > 1 ha, the total Landsat-measured pond reductions were similar to those derived using Sentinel-2. One advantage provided by the Landsat archive is an ability to examine beaver pond changes since 1985 compared to 2015 for Sentinel-2, which should better reveal any long-term cyclical patterns or associations with climate factors (Fraser et al. 2024). Overall, the consistent 2017-2023 beaver pond trends obtained using the three satellite sensors (Fig. 2, Fig. 4), and by comparing predictions from three-year Sentinel-2 composites to single images (Fig. 2b), increases confidence in the mapped beaver pond reductions.

Although the 2017-2023 beaver pond reductions were the largest observed during 1985-2023, two other periods (1986-1992 and 2001-2012) showed reductions that were up to 50% of this magnitude (Fig. 2a). No beaver census data were available for 1985-2014 to determine if this was related to earlier beaver population declines. However, these two periods coincided with the two lowest multiyear intervals of Lake Superior water levels since 1926, along with below average precipitation and SPEI values, suggesting that drought played a role. During 1985-2017, ponded area was most strongly correlated to Lake Superior water levels (Fig. 6a), which provided an effective integrated measure of regional water balance (Assel et al. 2004). The positive relationship between ponding and the climate variables then diverged after 2017 when their values remained above average and ponded area experienced the largest reduction since 1985.

The above observations highlight that potential drivers of ponding changes other than beaver occupancy must be considered if our satellite mapping approach is to provide a reliable indicator of beaver population trends. A major consideration would be long-term climate anomalies that may expand beaver-maintained ponds during wet years or shrink ponds during drought. In the case of Michipicoten Island, there was evidence that two periods of drought since 1985 caused relatively small (< 15%) beaver pond area reductions. In other regions, beaver ponding (Hood and Bayley 2008, Johnson-Bice 2021, Fitch et al. 2022) and lodge occupancy (Hood 2020) have not demonstrated strong relationships to climate, while beavers have been shown to buffer against the effects of drought (Hood and Bayley 2008, Fairfax and Small 2018). However, drought can cause short-term reductions in beaver pond depth (Hood and Bayley 2008, Naiman et al. 1994), as well as potentially decrease beaver colony density (Ribic et al. 2017) that could lead to further ponding reductions.

Although our study and others (Hood and Bayley 2008, Johnston and Windels 2015) show that a regional decline in beaver colonies should be detectable as a reduction in ponded area, the hydrologic response from an increasing beaver population may be more variable through time. Early regional beaver recolonization has been associated with a rapidly increasing pond and wetland area (Johnston and Naiman 1990, Snodgrass 1997, Cunningham et al. 2006, Martell et al. 2006, Hood and Bayley 2008, Johnston and Windels 2015, Martin et al. 2015, Johnson-Bice et al. 2022), but as beavers reach high densities, new colonies must occupy progressively less optimal habitats where smaller areas are impounded (Johnston and Naiman 1990, Cunningham et al. 2006, Johnson-Bice et al. 2022). Indeed, shortly after wolf colonization when beaver abundance was still very high (McLaren et al. 2022), field crews on Michipicoten Island observed beavers trying to impound water in virtually all small creeks, resulting in some “ponds” as small as 2-3 m^2^ (Patterson et al. pers. observation). This decreasing pond size at higher beaver densities may be offset by a residual or slowly decreasing surface water area in abandoned ponds that accumulate on the landscape through time (Johnston and Windels 2015, Johnson-Bice et al. 2022). Once a beaver population has become well established in a region, local ecological factors such as forage availability can create a dynamic mosaic of beaver patches that are periodically abandoned and drained, and recolonized and flooded (Meentemeyer and Butler 1995, Fryxell 2001, Kivinen et al 2020). This will lead to greater variability in populations (or ponding) at the local scale compared to the regional scale (Meentemeyer and Butler 1995, Fryxell 2001, Hartman et al. 2003, Johnson-Bice et al. 2022), especially in the absence of strong broad-scale drivers, such as climate extremes, epizootic disease, forage renewal after large wildfires (Fraser et al. 2025), or as in this study, new predation pressure in a closed system.

While our analysis shows that recent surface water area reductions on Michipicoten Island followed a large decline in beaver colonies, the probable ultimate cause for these changes was, at least in part, heavy beaver predation resulting from the arrival of grey wolves to the island over an ice bridge during the winter of 2013-2014 (McLaren et al. 2022). At that time, beavers were likely highly vulnerable to predation owing to their extremely dense population (6.1 colonies/km^2^) (Gable et al. 2018), which had expanded into shallow headwater creeks where relatively small ponds would offer less protection. Many beavers may also have had to travel progressively longer distances from the safety of their ponds in search of forage including during winter (Bergerud et al. 2020, Smith and Peterson 2021), where they would be more vulnerable in the presence of new predators (Gable et al. 2023). Although wintertime predation of beavers by wolves is considered rare (Hernandez and Bump 2022) it was a relatively common occurrence on Michipicoten Island and appeared to increase as caribou numbers declined following wolf colonization (Patterson et al. unpublished data). Further, scat analysis indicated that beavers were the most important summer food item for both adult wolves and their pups following their colonization on the island (Benson et al. 2021).

Beavers were also an important food source for wolves on Isle Royale, also in Lake Superior, where the 19 wolves translocated during 2018-2019 (8 being from Michipicoten Island) had a diet that was 62% beaver by frequency and 39% by biomass during the ice-free season (Sovie et al. 2023). Beavers had similarly reached historically high densities (1.0 colonies/km^2^) on Isle Royale following a previous wolf population collapse but after the wolf translocation, the number of colonies declined by 49% between 2021-2022 (Hoy et al. 2023).

This represents the third period since 1970 when the number of active beaver sites on Isle Royale has decreased in association with a rebounding wolf population (Smith and Peterson 2021), and we expect that would have led to beaver pond area reductions (Supplementary Video 3) as were mapped in this study. Overall, the strong response of beaver colony numbers to wolves on Michipicoten Island, Isle Royale, and the State Islands (Bergerud et al. 2020) in Lake Superior suggests that beaver populations in these closed systems have lower resilience to predation compared to other boreal systems where regional, source-sink metapopulation flow (Fryxell, 2001) may compensate for high predation rates (Gable and Windels 2017). We also show that on Michipicoten Island, predation likely resulted in larger and more persistent changes to total beaver ponded area than has been observed in northern Minnesota (Gable et al. 2020).

A limitation of the satellite-based pond mapping is that little or no surface water will be identified where there is a dense cover of emergent or floating aquatic vegetation (e.g. water lily and watershield) that produces a relatively high NIR reflectance. Therefore, if a beaver pond containing aquatic vegetation cover drains, reductions will be detected only in its open water portion (Fig. 5, Fig. S4). The impact of this would be to underpredict surface water declines in these ponds, producing a conservative estimate of beaver pond area reductions on Michipicoten Island. In addition, long-term shifts in aquatic vegetation cover resulting from changing grazing pressure (Bergman and Bump, 2015) or beaver pond maturation (Ray et al. 2001), would be reflected as surface water changes that may be unrelated to beaver occupancy or hydrologic conditions. To address this issue, we also tried using spring satellite composites (May 1 – June 15) for water mapping before most annual aquatic vegetation growth occurs. However, this led to new issues related to spring inundation of beaver pond meadows and other flat areas, and increased image noise when using a shorter spring compositing period.

Another consideration for our mapping method is that satellite reflectance was derived using three-year summer median values to ensure that input data were clean and free of atmospheric contamination. The resulting temporal smoothing could cause any transient, one-year ponding changes to go undetected. Nevertheless, the continuous satellite time series would be unlikely to miss multiyear, flooded vs drained state changes by comparison to the 5–10-year intervals of aerial photographs used in previous beaver pond mapping studies (Johnston and Naiman 1990, Meentemeyer and Butler 1995, Snodgrass 1997, Cunningham et al. 2006, Martell et al. 2006, Hood and Bayley 2008, Martin et al. 2015). Finally, for this approach to be applicable to analyzing larger regions, automated methods for identifying beaver pond complexes would have to be considered (Fairfax et al. 2023, Zhang et al. 2024) rather than relying on manual selection.

In summary, sub-pixel surface water mapping methods applied to Landsat and Sentinel-2 optical satellite data effectively measured a lagged reduction in beaver pond area after a large decline in the number of colonies on Michipicoten Island that was associated with the arrival of grey wolves. The largest changes occurred in smaller ponds, with Sentinel-2 mapping indicating a 48% decrease for the 73% of ponds that were < 0.1 ha. These reductions occurred during years with above average wetness conditions, but two smaller ponding reductions since 1985 were likely caused by drought. Beaver pond surface water changes mapped using medium resolution (10-30 m) satellite data may provide an efficient, cost-effective indicator of regional beaver population shifts and should be further tested. The approach may be especially suitable for vast, remote regions in the boreal zone where beaver ponds are relatively large, conventional beaver censuses are logistically prohibitive, and sub-meter commercial satellite or air photo imagery is too expensive or has limited availability.

## ACKNOWLEDGEMENTS

The authors would like to thank Gabriel Gosselin for orthorectifying high-resolution images, Morgan McFarlane-Winchester for assistance with GIS processing, Yu Zhang for extracting monthly and annual ERA5-Land precipitation data, and two anonymous reviewers of a previous version of this manuscript for their helpful comments and suggestions. Funding for this project was provided by the Earth Observation for Cumulative Effects project that is part of Natural Resources Canada’s Status and Trends Program.

## AUTHOR CONTRIBUTIONS

RF, IO, and AM conceived the ideas and designed methodology; RF, IO, AM, and BP collected the data; RF and IO analysed the data; RF led the writing of the manuscript. All authors contributed critically to the drafts and gave final approval for publication.

