## Supplementary Figures for "Beaver population decline on Michipicoten Island, Ontario leads to satellite-measured surface water area reductions"

Supplemental Figure 1. Comparison of beaver pond extents on Michipicoten Island (85.80°W, 47.75°N) visible in pan-sharpened WorldView-3 imagery (a) with maximum pond extents captured using (b) a Landsat binary surface water product (Pekel et al. 2016) and (c-d) Landsat and Sentinel-2 sub-pixel surface water products used in this study that estimate percentage water cover within each pixel.

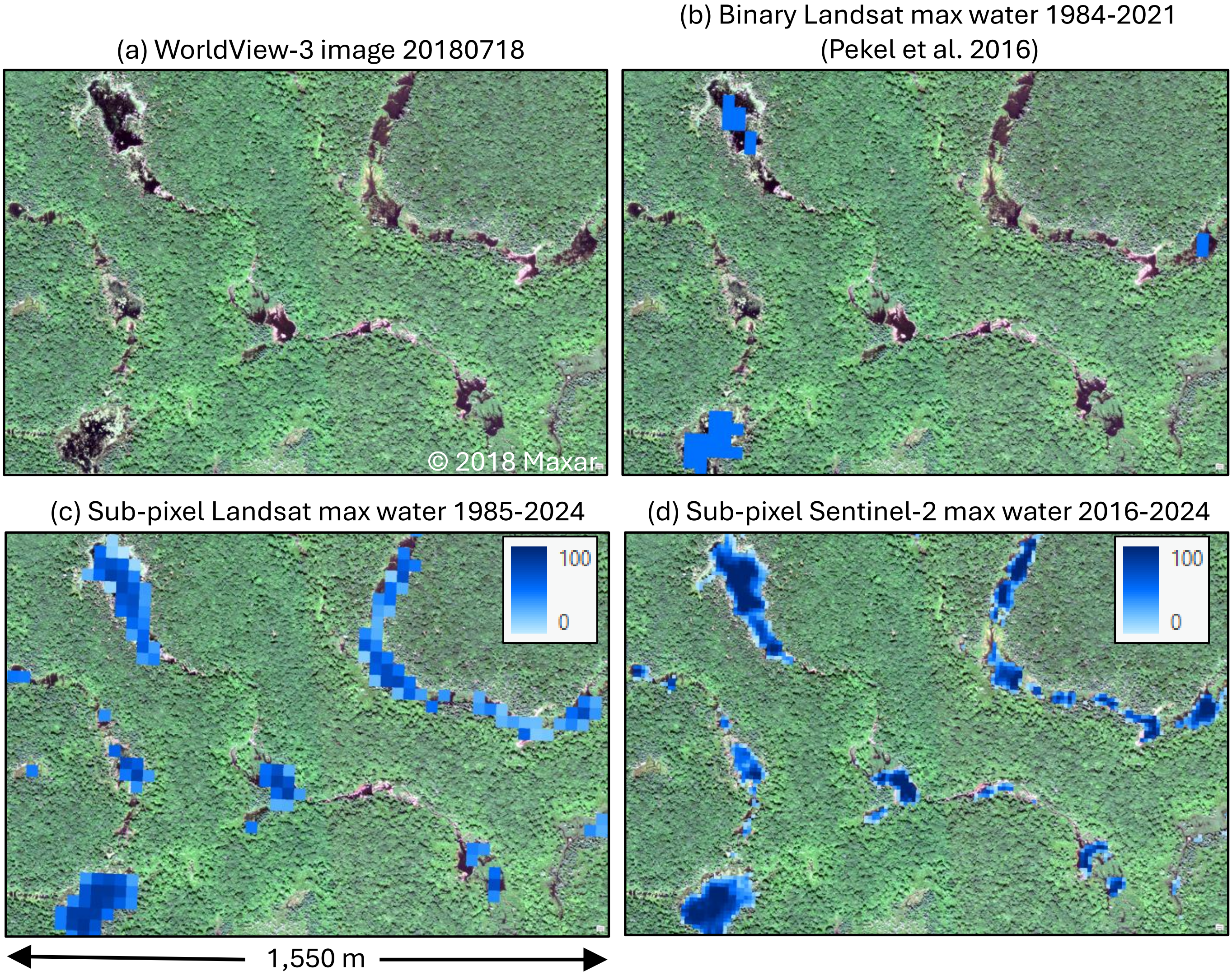



Pekel, J.F., Cottam, A., Gorelick, N., Belward, A., 2016. High-resolution mapping of global surface water and its long-term changes. Nature 540, 418–436. https://doi.org/10.1038/nature20584.

Supplemental Figure 2. Method used to mask needleleaf forest on Michipicoten Island, where erroneous, low satellite-based water fractions may be predicted. This commission error is more common in needleleaf forest recently impacted by likely insect defoliation (pink areas in the GeoEye-1 RGB image), which causes reductions in near-infrared reflectance that is used to map surface water area.

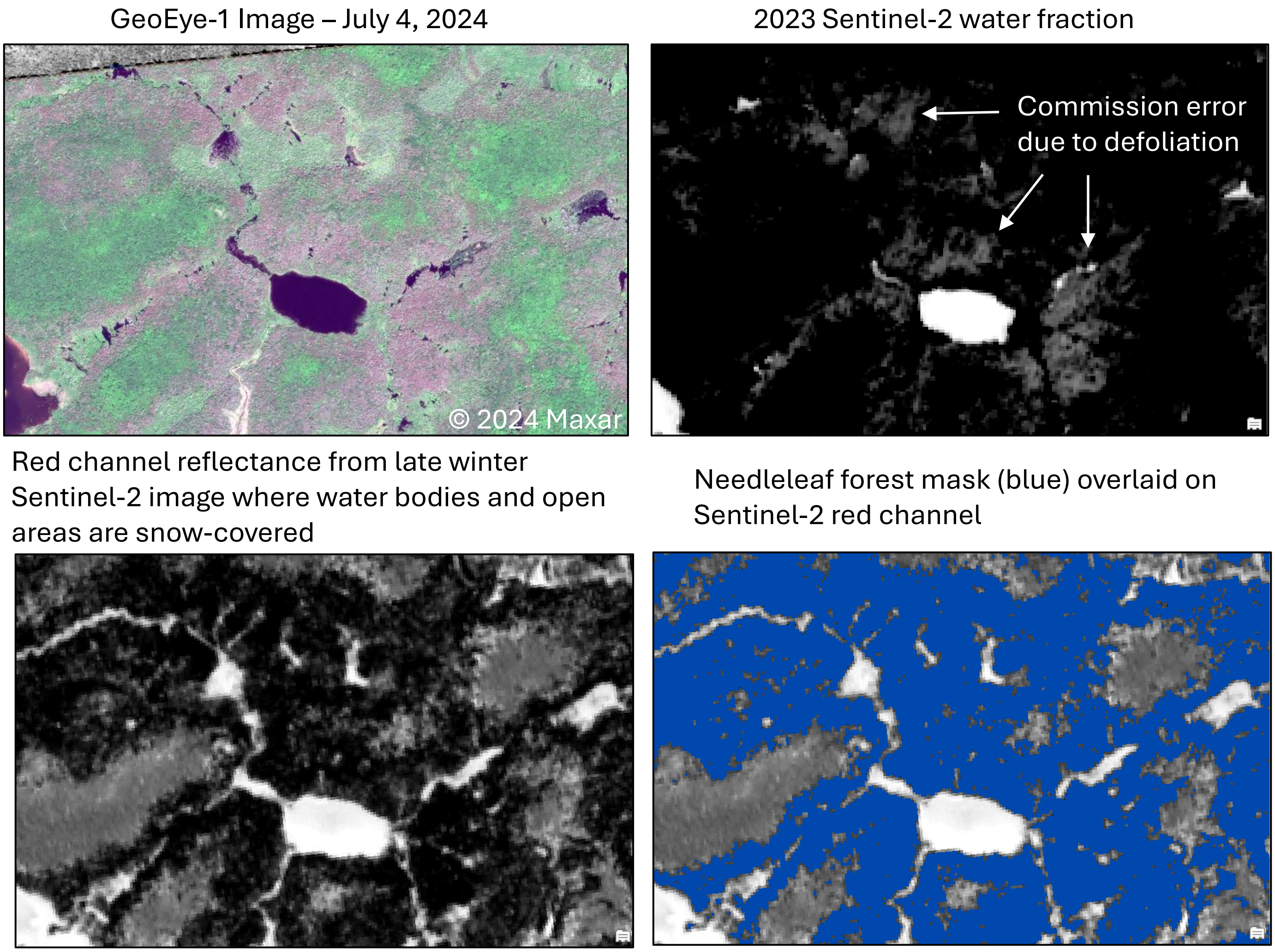

Supplemental Figure 3. (a) Example area (7.3 km$^2$) centred at 85.67°W, 47.75°N on Michipicoten Island showing pan-sharpened WorldView-3 imagery from July 18, 2018 (RGB=NIR,Green,Blue). All 170 water bodies mapped in this area using Sentinel-2 satellite imagery (dark blue polygons in (b)) are associated with visible beaver dams and in many cases beaver lodges and canals.

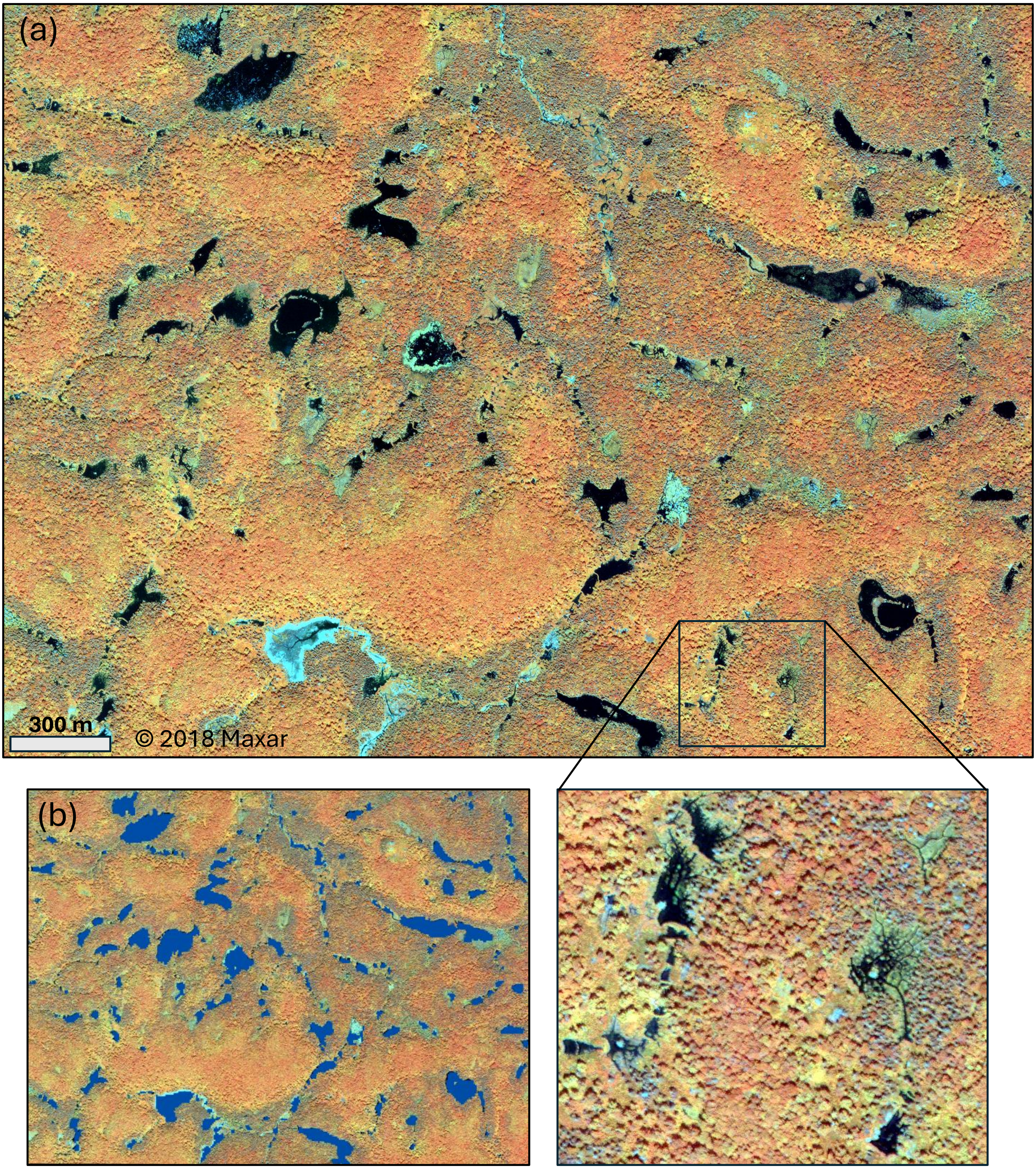

Supplemental Figure 4. A second example of beaver pond changes (2018-2024) mapped using high resolution WorldView-3 (a,c) and GeoEye-1 satellite images (b,d), and medium resolution Sentinel-2 (e-f) and Landsat (g-h) data. Surface water omission errors in 2018 can be observed at locations where there is a cover of floating aquatic vegetation.

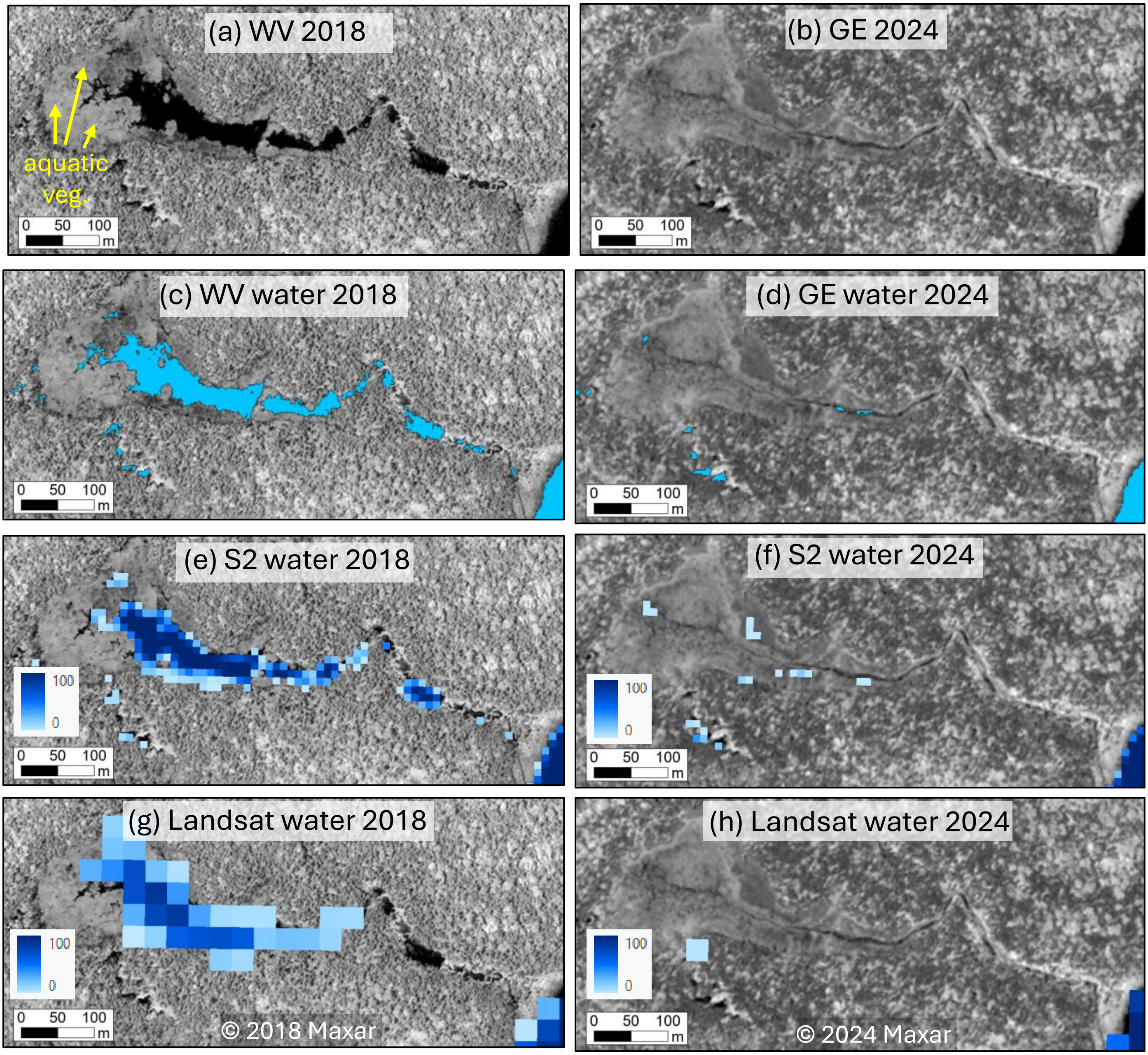
