## Supplementary Table for "Beaver population decline on Michipicoten Island, Ontario leads to satellite-measured surface water area reductions"

Supplementary Table 1. Date and Product ID for four clear-sky Level 1C Sentinel-2 images used to create single-date surface water maps on Michipicoten Island

| Image Date | Product ID |
|---|---|
| 20160705 | S2A_MSIL1C_20160705T164322_N0204_R126_T16TET_20160705T164320.SAFE |
| 20180725 | S2A_MSIL1C_20180725T163901_N0500_R126_T16TET_20230815T061423.SAFE |
| 20230818 | S2A_MSIL1C_20230818T163841_N0509_R126_T16TET_20230818T214646.SAFE |
| 20240718 | S2B_MSIL1C_20240718T163839_N0510_R126_T16TET_20240718T201929.SAFE |
