## Supplementary figures and images for "Beaver population decline on Michipicoten Island, Ontario leads to satellite-measured surface water area reductions"

### Supplementary Video 1

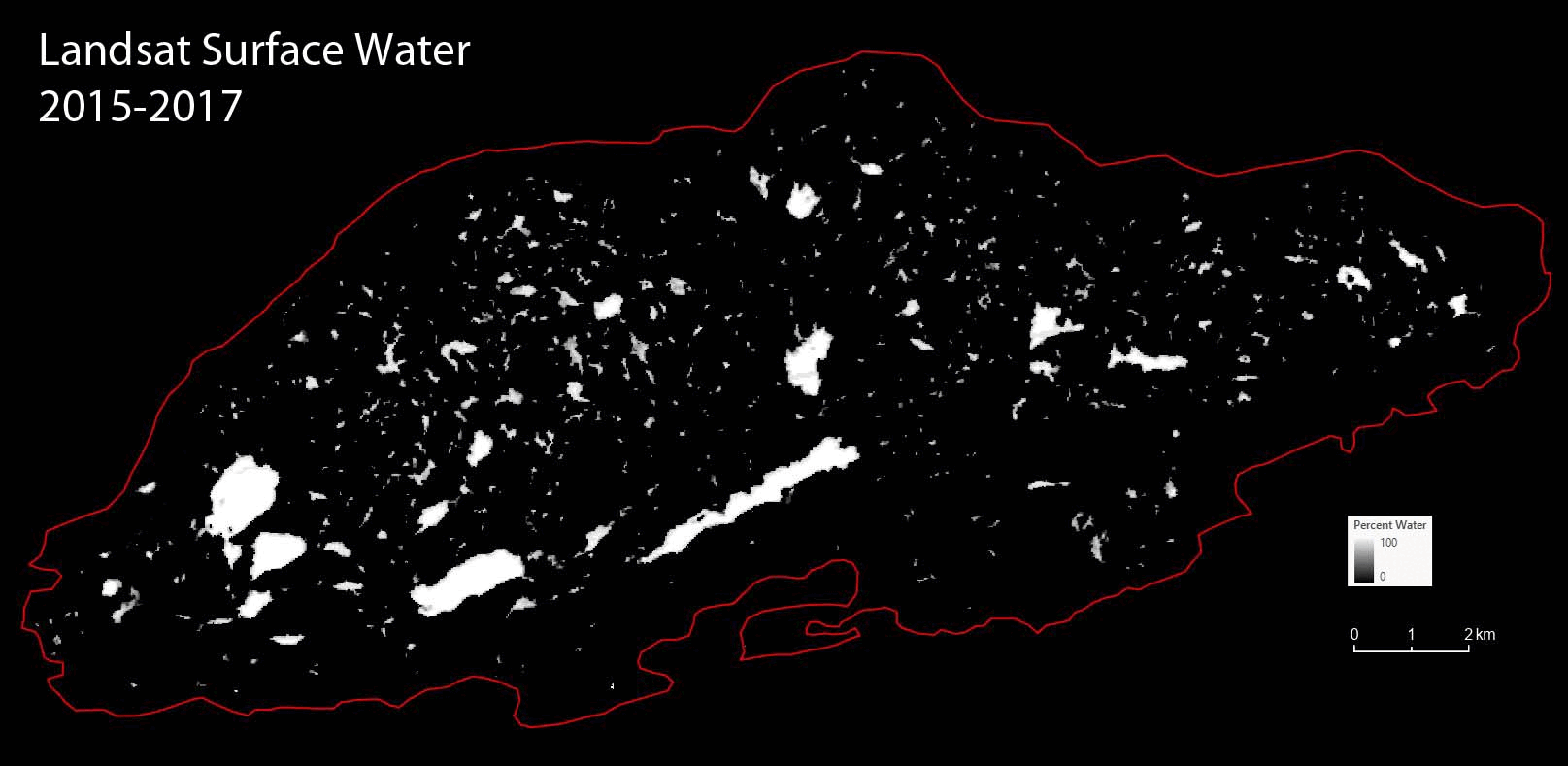

### Supplementary Video 2

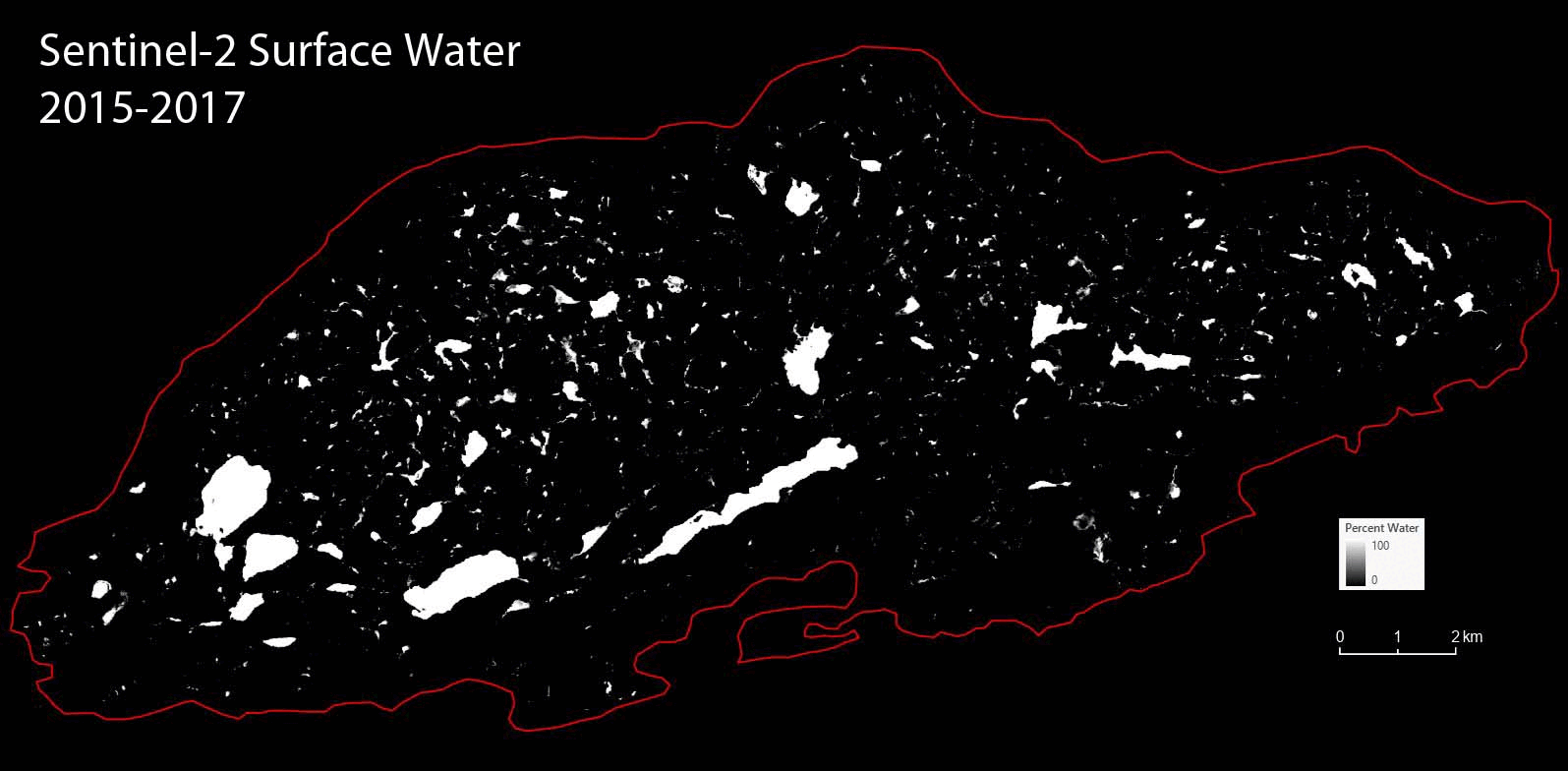

### Supplementary Video 3

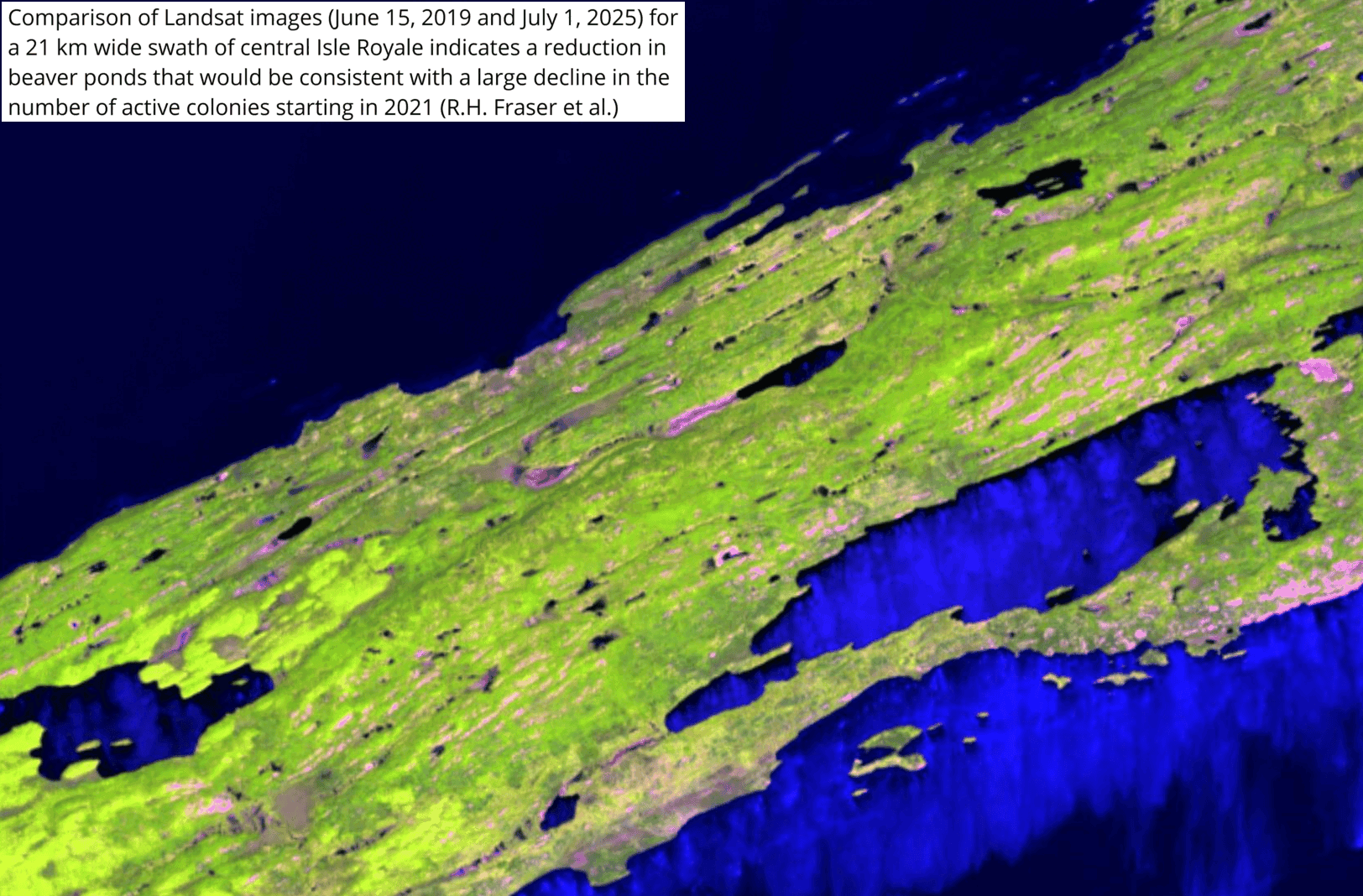
